# A new look at functional beta diversity

**DOI:** 10.1101/2024.02.25.580632

**Authors:** Carlo Ricotta, Sandrine Pavoine

## Abstract

1. The variability in species composition among a set of sampling sites, or beta diversity, is considered a key signature of the ecological processes that shape the spatial structure of species assemblages.
2. In this paper, we propose to decompose this variability into three additive components: i) the standard compositional similarity between individuals of the same species among sites, ii) the degree of functional *dissimilarity* between individuals of distinct species among sites, and iii) the degree of functional *similarity* between individuals of distinct species among sites. These three components can be used to portray the functional resemblance among sites on a ternary diagram.
3. The potential of this method is shown with real data on the functional turnover of Alpine species along a primary succession on glacial deposits in northern Italy.
4. ***Synthesis***. With the resulting ternary diagram of ‘functional resemblance’ we can relate various aspects of taxonomic and functional variability among sites to community assembly processes more completely than just looking at individual components.

## 1. Introduction

The amount of variation in species composition among sites, or beta diversity, is considered a fundamental tool for exploring the ecological processes that shape the spatial structure of species assemblages. Since the seminal work of Whittaker (1972), many different methods and measures have been proposed for summarizing beta diversity (Lande, 1996; Koleff et al., 2003; Anderson, 2006; Jost, 2007; Tuomisto, 2010a, 2010b; Anderson et al., 2011; Chao & Chiu, 2016; Legendre & De Caceres, 2013; Ricotta, 2017; Chao & Ricotta, 2019). One of the most commonly used consists in computing beta diversity as the mean compositional dissimilarity between pairs of sampling units (i.e., relevés, quadrats, assemblages, etc. which we will now generally refer to as plots). The general idea behind this approach is that for a set of plots, compositional heterogeneity or beta diversity increases with increasing mean dissimilarity between pairs of plots (Whittaker, 1972; Izsak & Price, 2001; Koleff et al., 2003; Chao & Chiu, 2016).

To compute dissimilarity-based beta diversity, standard measures, such as the Jaccard or the Bray-Curtis coefficients (Legendre & Legendre, 2012) were originally used. Such measures quantify taxonomic (i.e. species) differences between plots based only on species presences and absences or on abundance data, thus assuming that all species are equally and maximally distinct from each other, while neglecting information on functional differences among species. In the last decades however, several ‘functional dissimilarity measures’ have been proposed (reviewed in Lengyel & Botta-Dukát, 2023). Such measures take into account information on functional differences among species. Therefore, they are expected to improve correlation between community data and ecosystem functioning, as the species traits directly or indirectly influence these processes (Mouchet et al., 2010; Mason & de Bello, 2013).

A neglected outcome of the idea that distinct species possess varying degrees of functional dissimilarity (discussed by Ricotta et al., 2023 in the context of within-site diversity) is that the ecological information associated to the functional resemblance structure among plots is much richer and complex than that obtained from standard taxonomic dissimilarity measures.

Assuming that all species are equally and maximally distinct, standard similarity/dissimilarity measures in the range [0,1] are complementary to each other. For example, given two plots *h* and *k* with species relative abundances *p*_*jh*_ and *p* _*jk*_ (*j* =1, 2,…, *N*), the well-known Bray & Curtis (1957) dissimilarity and similarity coefficients *D*_*BC*_ and *S*_*BC*_ can be expressed as 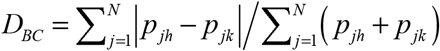 and 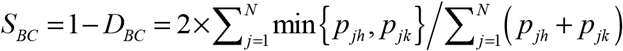, respectively, such that *S*_*BC*_ + *D*_*BC*_ = 1. Hence, looking only at one of them is enough to capture the entire information on the resemblance structure of both plots (the reason for using relative abundances instead of absolute abundances for the calculation of *D*_*BC*_ and *S*_*BC*_ will be clear in the following sections).

This is however not the case for functional resemblance, where the distinct species contribute to varying degrees to the similarity/dissimilarity structure among plots. In this latter case, it seems natural to decompose standard (abundance-based) taxonomic dissimilarity into two complementary functional components: the extent of functional *dissimilarity* among individuals of the species that differ between the two plots, and the extent of functional *similarity* among individuals of the species that differ between the two plots. For dissimilarity measures in the range [0,1] these two components, together with the standard taxonomic similarity between individuals of the *same* species in both plots, can be used to display the functional resemblance structure between plots on a ternary diagram.

An essential requirement to appropriately decompose functional resemblance is that, for a given pair of plots, functional dissimilarity is always lower than standard compositional dissimilarity. However, many of the existing measures of functional dissimilarity do not fulfill this requirement (Ricotta et al., 2020). In this paper, we will first introduce the proposed additive decomposition of functional beta diversity, together with its basic requirements. Next, the potential of this approach for a more comprehensive analysis of the amount of functional variation among sites is shown with a worked example on the functional turnover of Alpine species along a primary succession in northern Italy.

## 2. A step by step introduction to beta diversity decomposition

Let *d*_*ij*_ be a measure of functional dissimilarity between species *i* and *j* (*i, j* =1, 2,…, *N*) in the range [0,1] and *s*_*ij*_ be the corresponding functional similarity *s*_*ij*_ = 1− *d*_*ij*_. The functional dissimilarities *d*_*ij*_ summarize uni- or multivariate differences in the trait values between species such that *d*_*ij*_ = *d* _*ji*_ and *d*_*ii*_ = 0. For two plots *h* and *k*, let *D*_*F*_ be a measure of functional dissimilarity (0 ≤ *D*_*F*_ ≤ 1) that is computed by taking into account the actual functional differences *d*_*ij*_ between the species in both plots. Examples of such measures can be found e.g. in Rao (1982), Chao et al. (2014), Pavoine & Ricotta (2014), Ricotta et al. (2020, Appendix S1), Ricotta et al. (2021a) and in the worked example of this paper.

Further, let *D*_*S*_ be a corresponding measure of taxonomic dissimilarity between *h* and *k* (0 ≤ *DS* ≤ 1) that is computed solely from the differences in species abundances between both plots (i.e. assuming that all species are maximally dissimilar from each other, such that *d*_*ij*_ = 1 for all *i* ≠ *j*). If *D*_*S*_ ≥ *D*_*F*_, which is an intuitively reasonable condition given the definition of *D*_*F*_ and *D*_*S*_, we can decompose the resemblance structure between *h* and *k* into three additive components that describe distinct facets of the taxonomic and functional differences between both plots. For instance, the complements of *D*_*F*_ and *D*_*S*_ :

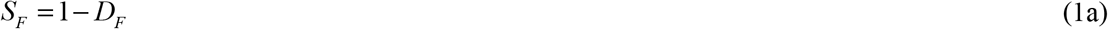

and

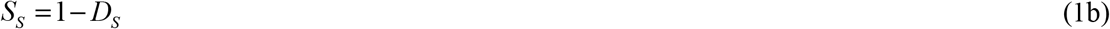

summarize the functional similarity *S*_*F*_ and the corresponding species similarity *S*_*S*_ (*S*_*S*_ ≤ *S*_*F*_) between the plots *h* and *k*, respectively. Like for *D*_*F*_ and *D*_*S*_, *S*_*F*_ is computed by taking into account the actual functional (dis)similarities between the species in both plots, whilst *S*_*S*_ is computed solely from the differences in species abundances between plots assuming that *s*_*ij*_ = 0 (*d*_*ij*_ = 1) for all species *i* ≠ *j*.

The third component of the proposed beta diversity decomposition is the difference between *D*_*S*_ and *D*_*F*_ (i.e. the fraction of species dissimilarity between *h* and *k* not expressed by functional dissimilarity):

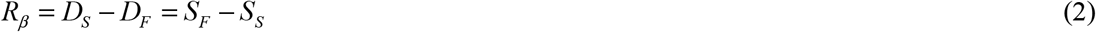

*R*_*β*_ represents the extent of functional *similarity* between individuals of the species unshared by the plots. From an ecological viewpoint, *R*_*β*_ can be interpreted as the degree to which individuals of species unshared by the plots support the same ecological functions. Therefore, Ricotta et al. (2020, 2021a) termed this quantity beta redundancy.

According to Eq. 2, species dissimilarity *D*_*S*_ can be thus additively decomposed into two distinct functional fractions: the degree of functional *dissimilarity* between individuals of distinct species *D*_*F*_ and the corresponding degree of functional *similarity* between individuals of distinct species, or beta redundancy *R*_*β*_ such that *D*_*S*_ = *D*_*F*_ + *R*_*β*_. Hence, the functional resemblance structure among plots is constituted of three distinct additive fractions, each with its own ecological meaning.

A relevant aspect of this decomposition is that for *d*_*ij*_ in the range [0,1] functional dissimilarity *D*_*F*_, beta redundancy *R*_*β*_, and species similarity *S*_*S*_ sum up to one:

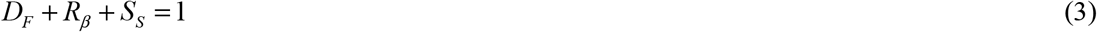

This offers the opportunity to use a ternary diagram to represent the functional resemblance structure among plots in graphical form. A ternary diagram displays the values of three variables *a, b*, and *c* as point coordinates on an equilateral triangle. The values of the variables must sum to a fixed constant, usually 1 (or 100%), such that *a* = 1− (*b* + *c*). The corners of the triangle represent a scenario in which one variable has a value of one and the other two variables have a value of zero. The values of each variable progressively decrease with increasing distance from the related corner (Ricotta et al., 2023). For example, if a point falls close to the *D*_*F*_ corner, this means that the corresponding pair of plots shows high functional dissimilarity, whereas closeness to the opposite side of the triangle reflects high functional similarity. Likewise, if a point falls close to the *R*_*β*_ corner, the corresponding pair of plots shows high beta redundancy; if the point falls close to the opposite side of this corner, the plots shows low beta redundancy.

With this ‘ternary diagram of functional resemblance’ we can thus graphically represent the compositional structure of a given set of plots in terms of pairwise functional dissimilarity, beta redundancy and species similarity. Therefore, ternary diagrams can be used to explore the ecological processes that shape different facets of the amount of variation in species composition among plots more exhaustively than by looking only at differences in functional dissimilarity (Podani & Schmera, 2011; Ricotta et al., 2023).

## 3. Worked example

### 3.1 Data

To illustrate the behavior of the proposed approach, we used the same data of Ricotta et al. (2021a, 2021b). The dataset is composed of a community composition matrix of 45 species in 59 plots of approximately 25 m^2^ sampled by Caccianiga et al. (2006) along a primary succession at the foreland of the Rutor Glacier (Northern Italy). The abundance of each species was assessed with a five-point ordinal scale transformed to ranks.

Based on the age of the glacial deposits, the plots were originally grouped by Caccianiga et al. (2006) into three successional stages. However, Ricotta et al. (2021a, 2021b) showed that in terms of functional beta diversity, the last two stages of the chronosequence are not significantly different from each other. Therefore, in this paper we classified all plots in the community composition matrix into two distinct groups: early successional plots (17 plots) and late successional plots (42 plots).

For all species, six key traits were used, which are related to the species global spectrum of form and function (Díaz et al., 2016): canopy height (CH; mm), leaf dry mass content (LDMC; %), leaf dry weight (LDW; mg), specific leaf area (SLA; mm^2^ × mg^−1^), leaf nitrogen content (LNC; %), and leaf carbon content (LCC; %). Data on species abundances and functional traits are available in Ricotta et al. (2016, Appendix S2), and Caccianiga et al. (2006, Table 2), respectively, and in the adiv (R package) object ‘`RutorGlacier`’ (Pavoine, 2020). The R scripts used in this paper are available in Appendix 1 of this paper.

### 3.2 Methods

The trait values were first standardized to zero mean and unit standard deviation. Next, like in Ricotta et al. (2021a, 2021b), we used the Euclidean distance to compute a matrix of pairwise functional distances between the 45 species from the standardized functional traits. The functional distances were then rescaled in the unit range by dividing each distance by the maximum value in the distance matrix. For all pair of plots in both successional stages, we finally used the algorithmic dissimilarity coefficient of Kosman (1996) and Gregorius et al. (2003) *D*_*KG*_ to calculate the three components of functional resemblance: functional dissimilarity, beta redundancy and species similarity.

A necessary condition to additively decompose functional resemblance is that, for two plots *h* and *k*, functional dissimilarity *D*_*F*_ is always lower than species dissimilarity *D*_*S*_. This prevents negative redundancy values, which would obviously be meaningless. However, as shown by Ricotta et al. (2020, Appendix S1), virtually none of the existing measures of functional dissimilarity conforms to this ‘redundancy property’.

Unlike most standard analytical measures of functional dissimilarity, the algorithmic dissimilarity *D*_*KG*_ conforms to the redundancy property (proof in Ricotta et al., 2021a, Appendix S1). The measure, which has been originally developed to measure genetic differences between populations, is based on the best possible match between the species in *h* and *k* in order to minimize the total functional differences between the plots. For two plots *h* and *k*, with *n* individuals in both plots, each individual in *h* is matched to an individual in *k* with the goal of minimizing the sum of functional differences between the individuals in both plots. The *n* pairs are formed such that all individuals in each plot are used only once. The functional dissimilarity *D*_*KG*_ is then obtained as the mean dissimilarity between each pair of individuals (Kosman & Leonard, 2007). However, since the actual number of individuals in *h* and *k* is usually not the same, to get a complete association between the individuals in both plots, the matching procedure is performed on the species relative abundances *p*_*jh*_ and *p*_*jk*_ in *h* and *k*, respectively.

Mathematically, the functional dissimilarity *D*_*KG*_ between plots *h* and *k* can be formulated as (Gregorius et al., 2003):

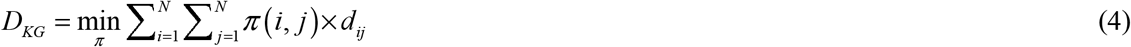

where *π* (*i, j*) is the relative abundance of species *i* in plot *h* that is matched with species *j* in plot *k*.

The use of species relative abundances *p* _*jh*_ (with 0 ≤ *p* _*jh*_ ≤ 1 and 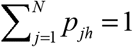) for the calculation of functional dissimilarity is justified by the observation that in most cases, ecologists are interested in exploring how the fraction of individuals that support a given ecological function differ between two plots (i.e. how the functional traits are proportionally distributed among the species in both plots), irrespective of the species absolute abundances in each plot.

Finding the optimal association between the species abundances in *h* and *k* is a special kind of linear optimization problem (Dantzig & Thapa, 1997). Since *D*_*KG*_ is essentially a mean dissimilarity between pairs of individuals, if the functional dissimilarity *d*_*ij*_ ranges from 0 (minimal dissimilarity between matched individuals) to 1 (maximal dissimilarity between matched individuals), *D*_*KG*_ also ranges from 0 to 1. Similarly, the complement of functional dissimilarity *D*_*KG*_ represents a measure of pairwise functional similarity *S*_*KG*_ = 1− *D*_*KG*_ between *h* and *k* that can be calculated as the optimal matching between the species abundances in *h* and *k* so as to *maximize* the mean similarity *s*_*ij*_ = 1 − *d*_*ij*_ between the species in both plots.

Kosman (2014) further showed that if all species in *h* and *k* are maximally dissimilar from each other (i.e. if *d*_*ij*_ =1 for all species *i* ≠ *j*), functional dissimilarity reduces to the classical Bray-Curtis dissimilarity computed from the species relative abundances *p* _*jk*_ :

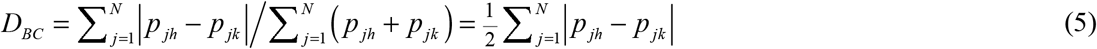

such that *D*_*KG*_ ≤ *D*_*BC*_. This allows to decompose functional similarity *S*_*KG*_ into standard taxonomic similarity between the individuals of the same species in both plots

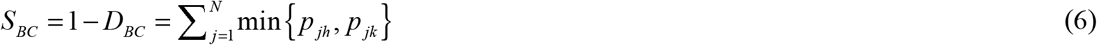

and the degree of functional similarity between the individuals of distinct species in both plots, or beta redundancy *R*_*β*_ = *D*_*BC*_ − *D*_*KG*_ = *S*_*KG*_ − *S*_*BC*_ such that *D*_*KG*_ + *R*_*β*_ + *S*_*BC*_ = 1 (see Eq. 3). An R function for the calculation of *D*_*BC*_, *D*_*KG*_ and *R*_*β*_ can be found in the R package adiv (Pavoine, 2020).

To test for differences in the beta diversity of both successional stages using all three components of the functional resemblance structure, we calculated the mean values of functional dissimilarity 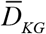, beta redundancy 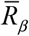, and species similarity 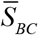 of each plot from all other plots of the same successional stage (see Ricotta et al., 2021b). The resulting values were then plotted on the ternary diagram of Figure 1 with the R package adegraphics (Siberchicot et al., 2017).

**Figure 1.**
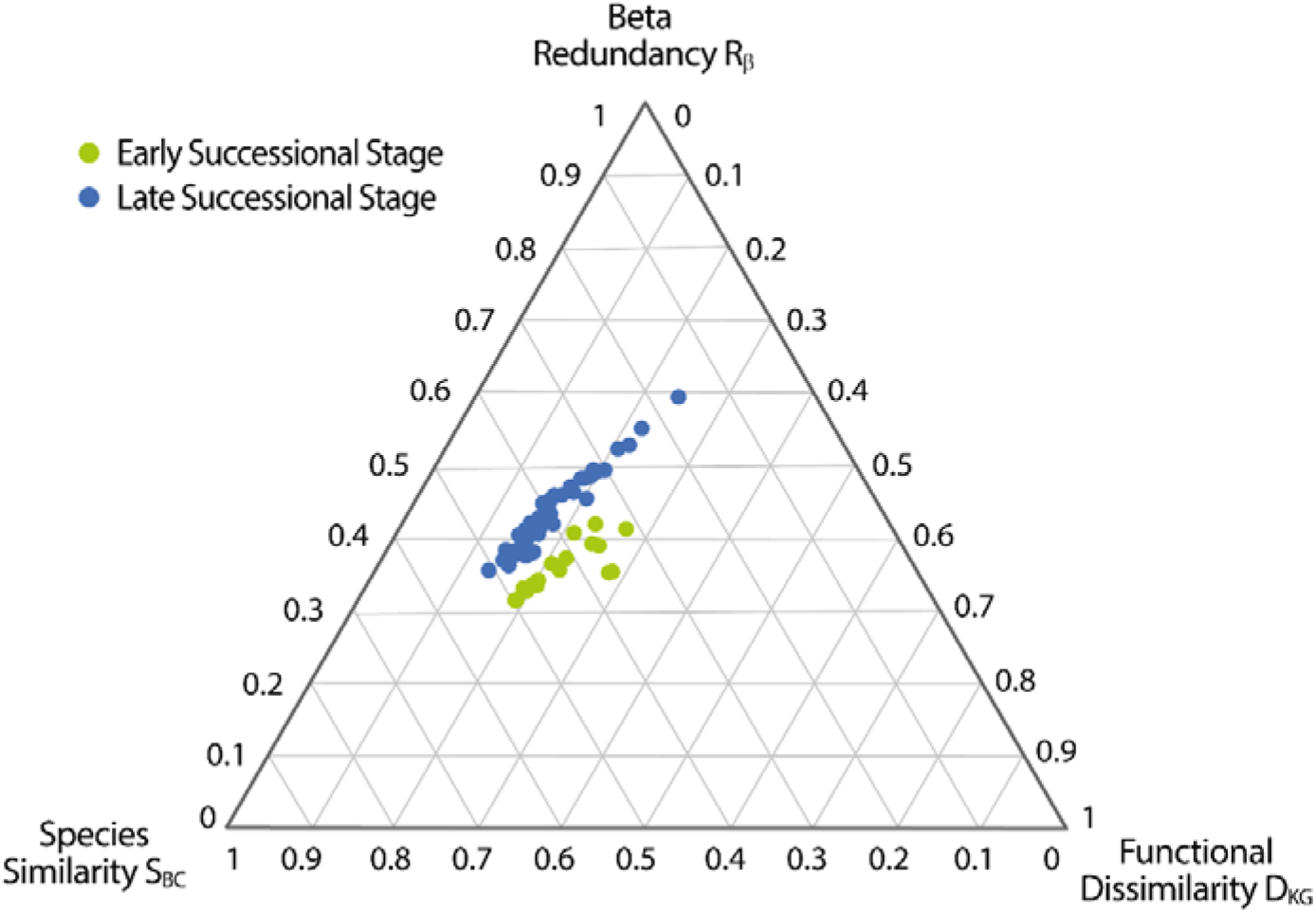
Ternary diagram of functional resemblance for the early and late successional plots of Alpine vegetation on glacial deposits in northern Italy. The results of db-MANOVA show that the two successional stages significantly differ in their overall functional resemblance structure at p < 0.001 (pseudo-F = 10.23, Bray-Curtis dissimilarity and 10000 randomizations).

Once the mean values 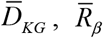, and 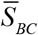 of the two groups of plots have been plotted on the ternary diagram, testing for differences in functional resemblance between the two successional stages of the Alpine vegetation essentially reduces to testing whether the distribution of the two groups of plots on the ternary diagram does not overlap. Therefore, we tested for differences in the ternary composition of both groups of plots with distance-based multivariate analysis of variance (db-MANOVA; Anderson, 2001) and the Bray-Curtis dissimilarity with the R package PERMANOVA (Vicente-Gonzalez & Vicente-Villardon, 2021).

db-MANOVA is a multivariate extension of standard analysis of variance which uses any multivariate dissimilarity measure of choice to test for differences between two or more distinct groups of plots (Anderson, 2001). The purpose of db-MANOVA is to contrast the within-group dissimilarities among plots with their between-group dissimilarities. The greater the between-group dissimilarities in comparison to the within-group dissimilarities, the more likely the groups of plots differ in their ternary composition (Anderson, 2001). P-values were calculated using 10000 permutations of single plots between the distinct groups. In this way, the single plots were randomly reassigned to the two successional stages while maintaining the vector of the functional resemblance values 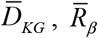 and 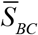 of each plot unchanged (Ricotta et al., 2021b).

At least for exploratory data analysis, Ricotta et al. (2023) considered this procedure appropriate for handling compositional data with a constant sum constraint. Those looking for statistical methods explicitly developed for the analysis of compositional data can refer to Aitchison (1986) or Van den Boogaart & Tolosana-Delgado (2013).

Finally, for each single resemblance measure, 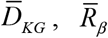 and 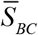, we separately tested for differences between the two successional stages with standard univariate ANOVA and 10000 permutations of individual observations between both groups of plots. Since db-MANOVA does not identify which particular resemblance measure is significantly different between groups of plots, the analysis of variance of the single components 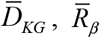 and 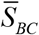 can then be used as a kind of post-hoc test to explore differences between multiple groups using each component at a time.

## 4. Results

Ternary diagrams have been first used for the analysis of beta diversity/dissimilarity by Podani & Schmera (2011). Such diagrams allow us to visualize the relative fractions of three variables on a two-dimensional graph. Aside from visual inspection, the point patterns in a ternary diagram can also be analyzed statistically. According to db-MANOVA, the successional stages in the ternary diagram of Figure 1 significantly differ in their overall functional resemblance structure (F = 10.23, p < 0.001). As shown in Table 1, the random dispersal mechanisms that drive the colonization of the glacial deposits in the early successional stages give rise to significantly higher values of mean functional dissimilarity between plots 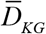, and to lower values of functional redundancy among individuals of different species 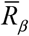 By contrast, while the species turnover among plots 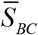 is approximately the same in both successional stages, in the late successional stages the species in one plot tend to be replaced by functionally similar species in the other plots, thus leading to an increased level of functional beta redundancy among different sampling units (Caccianiga et al., 2006).

**Table 1.** Mean (SD) values of average functional dissimilarity 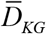, beta redundancy 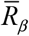 and species similarity 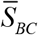 of each plot from all other plots of the same successional stage. Pairwise comparisons of index differences between both successional stages were performed with standard univariate ANOVA. P-values were obtained by randomly permuting individual plots between the successional stages (10000 permutations). *** = significant at p < 0.001; NS = not significant at p < 0.05.

|  | Early successional<br>plots (17 plots) | Late successional<br>plots (42 plots) |
| --- | --- | --- |
| Functional<br>dissimilarity $\bar{D}_{KG}$ *** | 0.217 (0.033) | 0.165 (0.024) |
| Beta<br>redundancy $\bar{R}_\beta$ *** | 0.370 (0.030) | 0.440 (0.050) |
| Species similarity<br>$\bar{S}_{BC}$ NS | 0.413 (0.054) | 0.395 (0.072) |

## 5. Discussion

The aim of this paper was to bring together distinct aspects of the analysis of functional beta diversity, redundancy, and community similarity into a coherent system. Some of these results were previously discussed in the context of within-site diversity, or alpha diversity (e.g. Ricotta et al., 2023) but can easily be extended to beta diversity. Resemblance measures that can be partitioned into complementary components are extremely valuable since the resultant elements can be related to a variety of distinct ecological processes that determine the structure of species assemblages (Baselga, 2010; Podani & Schmera, 2011; Podani et al., 2013; Ricotta et al., 2023). In this context, the relevant questions are: how to decompose the pairwise functional resemblance between plots, how to graphically represent this decomposition and how to test for significant differences in functional resemblance among groups of plots.

Unlike standard similarity/dissimilarity coefficients which possess a simple binary structure (the individuals in both plots are either totally similar to each other if they belong to the same species, or totally distinct if they belong to different species), functional resemblance can be additively decomposed into three complementary components: functional dissimilarity *D*_*F*_, beta redundancy *R*_*β*_ and species similarity *S*_*S*_. This increased complexity arises from the relaxation of the constraint that all species are equally and maximally distinct. For example, if different species possess varying degrees of functional dissimilarity, two plots with no species in common can be either functionally identical or completely distinct depending on the degree of functional dissimilarity *d*_*ij*_ between the species in both plots. Accordingly, looking only at differences in functional dissimilarity between plots provides just a partial view of the amount of taxonomic and functional variability between plots. To get a more comprehensive picture of the different facets of the resemblance structure between plots, the extent of functional similarity between individuals of distinct species needs to be also considered.

The concept of beta redundancy has been first introduced by Ricotta et al. (2020) as the amount of species dissimilarity between two plots not expressed by functional dissimilarity. Ricotta et al. (2020) used a relative measure of beta redundancy 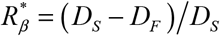. By contrast, due to the additive nature of the proposed dissimilarity decomposition, in this paper we used the absolute difference *R*_*β*_ = *D*_*S*_ − *D*_*F*_. Redundancy is maximal for two functionally identical plots with no species in common where *D*_*S*_ = 1 and *D*_*F*_ = 0. On the other hand, redundancy is zero when the species turnover between two plots is associated to a complete functional turnover such that *D*_*S*_ = *D*_*F*_ and hence *R*_*β*_ = 0.

Beta redundancy tells us to what degree the species turnover between two plots is associated to a functional turnover or, in other words, to what degree the species that differ between two plots are able to perform the same ecological functions. Accordingly, beta redundancy might be related to chief ecological processes, such as species dispersal, habitat filtering, or competitive exclusion. For instance, looking at the worked example, it can be assumed that since the values of mean species similarity among plots in both successional stages are more or less the same, the species dispersal ability did not change along the succession. By contrast, the lower functional dissimilarity and the higher beta redundancy of the late successional plots may be associated to the role of local species interactions (biotic filters) which constrain plant species recruitment more intensely compared to the abiotic filters of the early successional stages (Klanderud et al., 2010; Meineri et al., 2020).

From a more ‘technical’ viewpoint, in order to get a valid measure of beta redundancy, the functional dissimilarity *D*_*F*_ should always be lower than the corresponding species dissimilarity *D*_*S*_. Surprisingly, Ricotta et al. (2020) showed that many of the existing indices of functional dissimilarity do not conform to this ‘redundancy property’. Therefore, they cannot be used to appropriately decompose functional resemblance into non-negative additive fractions. In this view, the transition from a species-based ecology to a trait-based ecology has important implications not only in biological terms (Díaz & Cabido, 2001), but also in statistical terms.

To address this issue, Ricotta et al. (2020) introduced a tree-based measure of functional dissimilarity that conforms to the redundancy property. However, being based on a hierarchical representation which is the more natural way for describing the evolutionary relationships among species, the functional dissimilarity coefficient of Ricotta et al. (2020) is more adequate to represent the phylogenetic dissimilarity among plots rather than their functional differences. Ricotta et al. (2021a) thus suggested to summarize functional dissimilarity with the algorithmic coefficient of Kosman and Gregorius which does not depend on a tree-based species organization. While *D*_*KG*_ allows to calculate functional dissimilarity in accordance with the non-hierarchical nature of the species functional relationships, it does not exhaust the possibilities of measuring functional dissimilarity in an appropriate way. On the contrary, the search for a new class of functional dissimilarity measures that conform to the redundancy property seems a very promising research direction in order to enrich the ecologist’s toolbox with new, more up-to-date instruments to explore various aspects of functional resemblance and their ecological drivers.

## Supporting information

Appendix 1 - R scripts

## Author contributions

Carlo Ricotta conceptualized the research and wrote the original manuscript draft. Carlo Ricotta and Sandrine Pavoine analysed the data. Sandrine Pavoine wrote the R code to compute the multivariate dissimilarities among sites and to perform the beta diversity decomposition. Both authors contributed to editing the manuscript.

## Data availability Statement

Data on species abundances and functional traits are available in the adiv (R package) object ‘`RutorGlacier`’ (Pavoine, 2020). The R code used in this paper is available in Appendix 1 of this paper.

