## Appendix 1 - R scripts for "A new look at functional beta diversity"

**R scripts** used for the worked example of the functional beta diversity decomposition proposed in the main text. The scripts were written with R version 4.3.2.

**Disclaimer:** users of this code are cautioned that, while due care has been taken and it is believed accurate, its use and results are solely the responsibilities of the user.

```
# Package loading:

install.packages("adiv")
install.packages("adegraphics")
install.packages("PERMANOVA")

library(adiv) # version 2.2
library(adegraphics)# version 1.0-21
library("PERMANOVA") # version 0.2.0

# Data loading:

data(RutorGlacier)

# Functional dissimilarities between species:

fundis <- dist(scale(RutorGlacier$Traits2[1:6]))
fundis <- fundis/max(fundis)

# Relative abundance of species in plots:

prop <- sweep(RutorGlacier$Abund, 1, rowSums(RutorGlacier$Abund), "/")

# vector that indicates which plot belongs to which group (either early or late successional stage). Plots are in the same order as in table named prop above.

groups <- RutorGlacier$Fac
groups[groups == "mid"] <- "late"

# Data analyses:

propsplitted <- split(prop, as.factor(groups))
prop_Early <- propsplitted[[1]]
prop_Late <- propsplitted[[2]]

frameDKG_Early<- betaUniqueness(prop_Early, fundis)
frameDKG_Late <- betaUniqueness(prop_Late, fundis)

D_KG_Early<- frameDKG_Early$DKG  # Pairwise functional dissimilarities between plots of early successional stage.
S_BC_Early<- 1-frameDKG_Early$DR # Pairwise species similarity between plots of early successional stage.
R_beta_Early <- frameDKG_Early $DR- frameDKG_Early$DKG # Pairwise beta redundancy between plots of early successional stage.

D_KG_Late <- frameDKG_Late$DKG # Pairwise functional dissimilarities between plots of late successional stage.
S_BC_Late<- 1-frameDKG_Late$DR # Pairwise species similarity between plots of late successional stage.
R_beta_Late <- frameDKG_Late $DR- frameDKG_Late$DKG # Pairwise beta redundancy between plots of late successional stage.

# Below are calculated the average functional dissimilarities, beta redundancy and species similarity of each plot from the other plots of the same group. Plots are in the same order as in vector named "groups" and table named "prop".

D_KG_bar_Early <- sapply(1:17, function(i) mean(D_KG_Early[i, -i]))
S_BC_bar_Early<- sapply(1:17, function(i) mean(S_BC_Early[i, -i]))
R_beta_bar_Early <- sapply(1:17, function(i) mean(R_beta_Early[i, -i]))
D_KG_bar_Late <- sapply(1:42, function(i) mean(D_KG_Late[i, -i]))
S_BC_bar_Late<- sapply(1:42, function(i) mean(S_BC_Late[i, -i]))
R_beta_bar_Late <- sapply(1:42, function(i) mean(R_beta_Late[i, -i]))

# Graphical display for an equivalent of Figure 1 of main text

TAB_Early <- cbind.data.frame(D_KG_bar_Early, S_BC_bar_Early, R_beta_bar_Early)
TAB_Late <- cbind.data.frame(D_KG_bar_Late, S_BC_bar_Late, R_beta_bar_Late)
names(TAB_Early) <- names(TAB_Late) <- c("D_KG", "S_BC", "R_beta")
TAB <- rbind.data.frame(TAB_Early, TAB_Late)

triangle.class(TAB, as.factor(groups), starSize = 0, ellipseSize=0, adjust=FALSE, showposition =FALSE, col=c("green", "blue"))
```

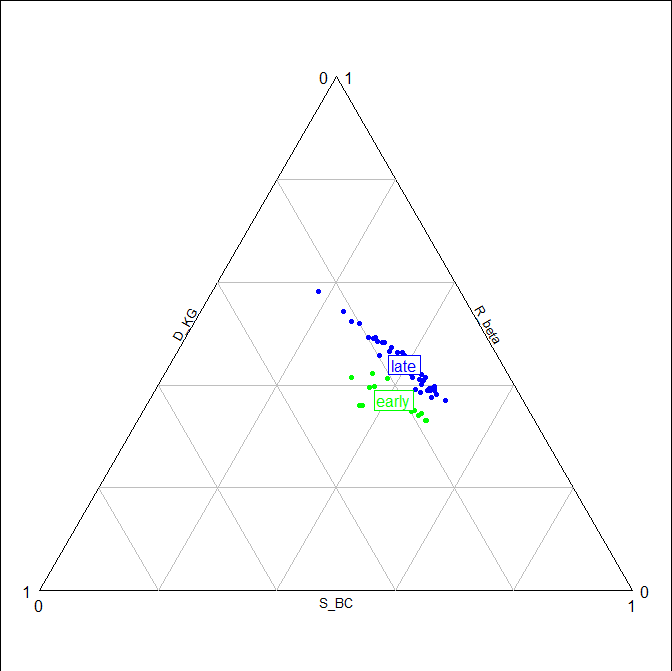

```
# Pairwise comparisons of index differences between both successional stages (Table 1 of main text)


# 1. Functional dissimilarity

mean(D_KG_bar_Early)
# [1] 0.2171854

sd(D_KG_bar_Early)
# [1] 0.03259282

mean(D_KG_bar_Late)
# [1] 0.164982

sd(D_KG_bar_Late)
# [1] 0.02423698


# 2. Beta redundancy

mean(R_beta_bar_Early)
# [1] 0.3701257

sd(R_beta_bar_Early)
# [1] 0.02989589

mean(R_beta_bar_Late)
# [1] 0.4396643

sd(R_beta_bar_Late)
# [1] 0.04991988

# 3. Species (dis)similarity

mean(S_BC_bar_Early)
# [1] 0.412689
sd(S_BC_bar_Early)
# [1] 0.0541672

mean(S_BC_bar_Late)
# [1] 0.3953538

sd(S_BC_bar_Late)
# [1] 0.07227821

# 4. Global PERMANOVA test

TAB4Ptest <- DistContinuous(TAB, , "Bray_Curtis")

Ptest <- PERMANOVA(TAB4Ptest, as.factor(groups), nperm=10000)

Ptest
###### PERMANOVA Analysis #######
# MANOVA
#      Explained  Residual df Num df Denom    F-exp    p-value p-value adj.
# Total 0.047524   0.26475      1       57 10.23187 0.00089991   0.0008999
```
